# *Candidatus* Ethanoperedens, a thermophilic genus of archaea mediating the anaerobic oxidation of ethane

**DOI:** 10.1101/2020.03.21.999862

**Authors:** Cedric Jasper Hahn, Rafael Laso-Pérez, Francesca Vulcano, Konstantinos-Marios Vaziourakis, Runar Stokke, Ida Helene Steen, Andreas Teske, Antje Boetius, Manuel Liebeke, Rudolf Amann, Katrin Knittel, Gunter Wegener

## Abstract

Cold seeps and hydrothermal vents deliver large amounts of methane and other gaseous alkanes into marine surface sediments. Consortia of archaea and partner bacteria thrive on the oxidation of these alkanes and its coupling to sulfate reduction. The inherently slow growth of the involved organisms and the lack of pure cultures have impeded the understanding of the molecular mechanisms of archaeal alkane degradation. Here, using hydrothermal sediments of the Guaymas Basin (Gulf of California) and ethane as substrate we cultured microbial consortia of a novel anaerobic ethane oxidizer *Candidatus* Ethanoperedens thermophilum (GoM-Arc1 clade) and its partner bacterium *Candidatus* Desulfofervidus auxilii previously known from methane-oxidizing consortia. The sulfate reduction activity of the culture doubled within one week, indicating a much faster growth than in any other alkane-oxidizing archaea described before. The dominance of a single archaeal phylotype in this culture allowed retrieving a closed genome of *Ca*. Ethanoperedens, a sister genus of the recently reported ethane oxidizer *Candidatus* Argoarchaeum. The metagenome-assembled genome of *Ca*. Ethanoperedens encoded for a complete methanogenesis pathway including a methyl-coenzyme M reductase (MCR) that is highly divergent from those of methanogens and methanotrophs. Combined substrate and metabolite analysis showed ethane as sole growth substrate and production of ethyl-coenzyme M as activation product. Stable isotope probing showed that the enzymatic mechanisms of ethane oxidation in *Ca*. Ethanoperedens is fully reversible, thus its enzymatic machinery has potential for the biotechnological development of microbial ethane production from carbon dioxide.

**IMPORTANCE:** In the seabed gaseous alkanes are oxidized by syntrophic microbial consortia that thereby reduce fluxes of these compounds into the water column. Because of the immense quantities of seabed alkane fluxes, these consortia are key catalysts of the global carbon cycle. Due to their obligate syntrophic lifestyle, the physiology of alkane-degrading archaea remains poorly understood. We have now cultivated a thermophilic, relatively fast-growing ethane oxidizer in partnership with a sulfate-reducing bacterium known to aid in methane oxidation, and have retrieved the first complete genome of a short-chain alkane-degrading archaeon. This will greatly enhance the understanding of non-methane alkane activation by non-canonical methyl-coenzyme M reductase enzymes, and provide insights into additional metabolic steps and the mechanisms underlying syntrophic partnerships. Ultimately, this knowledge could lead to the biotechnological development of alkanogenic microorganisms to support the carbon neutrality of industrial processes.

**Etymology:** *Ethanoperedens. ethano*, (new Latin): pertaining to ethane; *peredens* (Latin): consuming, devouring; *thermophilum*. (Greek): heat-loving. The name implies an organism capable of ethane oxidation at elevated temperatures.

**Locality:** Enriched from hydrothermally heated, hydrocarbon-rich marine sediment of the Guaymas Basin at 2000 m water depth, Gulf of California, Mexico.

**Diagnosis:** Anaerobic, ethane-oxidizing archaeon, mostly coccoid, about 0.7 μm in diameter, forms large irregular cluster in large dual-species consortia with the sulfate-reducing partner bacterium ‘*Candidatus* Desulfofervidus auxilii’.

## INTRODUCTION

In deep marine sediments, organic matter undergoes thermocatalytic decay, resulting in the formation of natural gas (methane to butane) and crude oil. If not capped, the gas fraction will rise towards the sediment surface due to buoyancy, porewater discharge and diffusion. Most of the gas is oxidized within the sediments coupled to the reduction of the abundant electron acceptor sulfate [1, 2]. Responsible for the anaerobic oxidation of alkanes are either free-living bacteria or microbial consortia of archaea and bacteria. Most free-living bacteria use alkyl succinate synthases to activate the alkane, forming succinate-bound alkyl units as primary intermediates [3]. Usually, these alkanes are completely oxidized, and this process is coupled to sulfate reduction in the same cells, as has been shown, for example in the deltaproteobacterial butane-degrading strain BuS5 [4]. However, alkane oxidation in seafloor sediments is too a large extent performed by dual species consortia of archaea and bacteria [5, 6]. As close relatives of methanogens, the archaea in those consortia activate alkanes as thioethers and completely oxidize the substrates to CO_2_. The electrons released during alkane oxidation are consumed by the sulfate-reducing partner bacteria.

The anaerobic methane-oxidizing archaea (ANME) activate methane using methyl-coenzyme M reductases (MCR) that are highly similar to those of methanogens, forming methyl-coenzyme M as primary intermediate [7]. The methyl group is oxidized via a reversal of the methanogenesis pathway [8]. Thermophilic archaea of the genus *Candidatus* Syntrophoarchaeum thrive on the oxidation of butane and propane. In contrast to ANME, they contain four highly divergent MCR variants, which generate butyl- and propyl-coenzyme M (CoM) as primary intermediates [9]. Based on genomic and transcriptomic evidence the CoM-bound alkyl units are transformed to fatty acids and oxidized further via beta-oxidation. The reactions transforming the CoM-bound alkyl units to CoA-bound fatty acids and the enzymes performing such reactions are so far unknown. The CoA-bound acetyl units are completely oxidized in the Wood-Ljungdahl pathway including the upstream part of the methanogenesis pathway. In hydrogenotrophic methanogens, the enzymes of this pathway are used to reduce CO_2_-forming methyl-tetrahydromethanopterin for methanogenesis and for biomass production. In *Ca*. Syntrophoarchaeum this pathway is used in reverse direction for the complete oxidation of acetyl-CoA. Both, the thermophilic ANME-1 and Ca. Syntrophoarchaeum form dense consortia with their sulfate-reducing partner bacterium *Candidatus* Desulfofervidus (HotSeep-1 clade) [10, 11]. The transfer of reducing equivalents between the alkane oxidizing archaea and their partners is likely mediated by pili-based nanowires and cytochromes produced by both consortial partners [12]. For a critical view on electron transfer in AOM consortia see [13].

Sulfate-dependent ethane oxidation has been described multiple times in slurries of marine sediments [4, 14, 15]. The first functional description of this process based on a cold-adapted culture derived from Gulf of Mexico sediments [5]. In this culture, *Candidatus* Argoarchaeum (formerly known as GoM-Arc1 clade) activates ethane with the help of divergent MCRs that are phylogenetically placed on a distinct branch next to those of *Ca*. Syntrophoarchaeum. Based on the presence of all enzymes of the Wood-Ljungdahl pathway that can be used for acetyl-CoA oxidation, it has been suggested that the CoM-bound ethyl groups are transferred to CoA-bound acetyl units. The required intermediates for this reaction mechanism are so far unknown [5]. *Ca*. Argoarchaeum forms unstructured consortia with yet unidentified bacterial partners and grows slowly with substrate turnover rates comparable to AOM [5]. Additional metagenome assembled genomes (MAGs) of the GoM-Arc1 clade derived from the Guaymas Basin and the Gulf of Mexico have similar gene contents, suggesting that these GoM-Arc1 archaea are ethane oxidizers [16, 17].

To date, the understanding of short-chain alkane metabolizing archaea mainly relies on comparison of their genomic information with those of methanogens that are well-characterized with regard to their enzymes. Due to the slow growth of the alkane-oxidizing archaea and the resulting lack of sufficient biomass, specific biochemical traits remain unknown. For instance, the structural modifications of non-canonical MCRs, or the proposed transformation of the CoM-bound alkyl to CoA-bound acetyl units in the short-chain alkane degraders has not been proven. Here, we describe a faster growing, thermophilic ethane-oxidizing culture from sediments of the Guaymas Basin. Metagenomic analyses of Guaymas Basin sediments revealed a great diversity of potential alkane degraders with divergent MCR enzymes [9, 18]. With ethane as sole energy source and sulfate as electron acceptor we obtained well-growing meso- and thermophilic ethane-degrading enrichment cultures from these sediments. Their low strain diversity makes them particularly suitable for assessing the pathways of the anaerobic oxidation of ethane.

## RESULTS AND DISCUSSION

### Establishment of meso- and thermophilic ethane-oxidizing enrichment cultures

Sediments were sampled from the gas- and oil-rich sediments covered by sulfur-oxidizing mats of the Guaymas Basin. From these sediments and artificial seawater medium, a slurry was produced under anoxic conditions and distributed into replicate bottles. These bottles were supplied with an ethane headspace (2 atm), and incubated at 37°C and 50°C. Additional growth experiments were performed with methane and controls were set up with a nitrogen atmosphere. As measure of metabolic activity, sulfide concentrations were tracked over time (further details see methods). Both methane and ethane additions resulted in the formation of 15 mM sulfide within 4 months. Instead, nitrogen controls produced only little sulfide (< 2mM) that likely corresponds to the degradation of alkanes and organic matter from the original sediment. Subsequent dilution (1:3) of the ethane and methane cultures and further incubation with the corresponding substrates showed faster, exponentially increasing sulfide production in the ethane culture, suggesting robust growth of the ethane-degrading community (Fig. 1 A). After three consecutive dilution steps, virtually sediment-free cultures were obtained. These cultures produced approximately 10 mM sulfide in 8 weeks. All further experiments were conducted with the faster growing 50°C culture (Ethane50). Sequencing of metagenomes, however, was done on both, the 50°C and 37°C (Ethane37) culture.

**Fig. 1.**
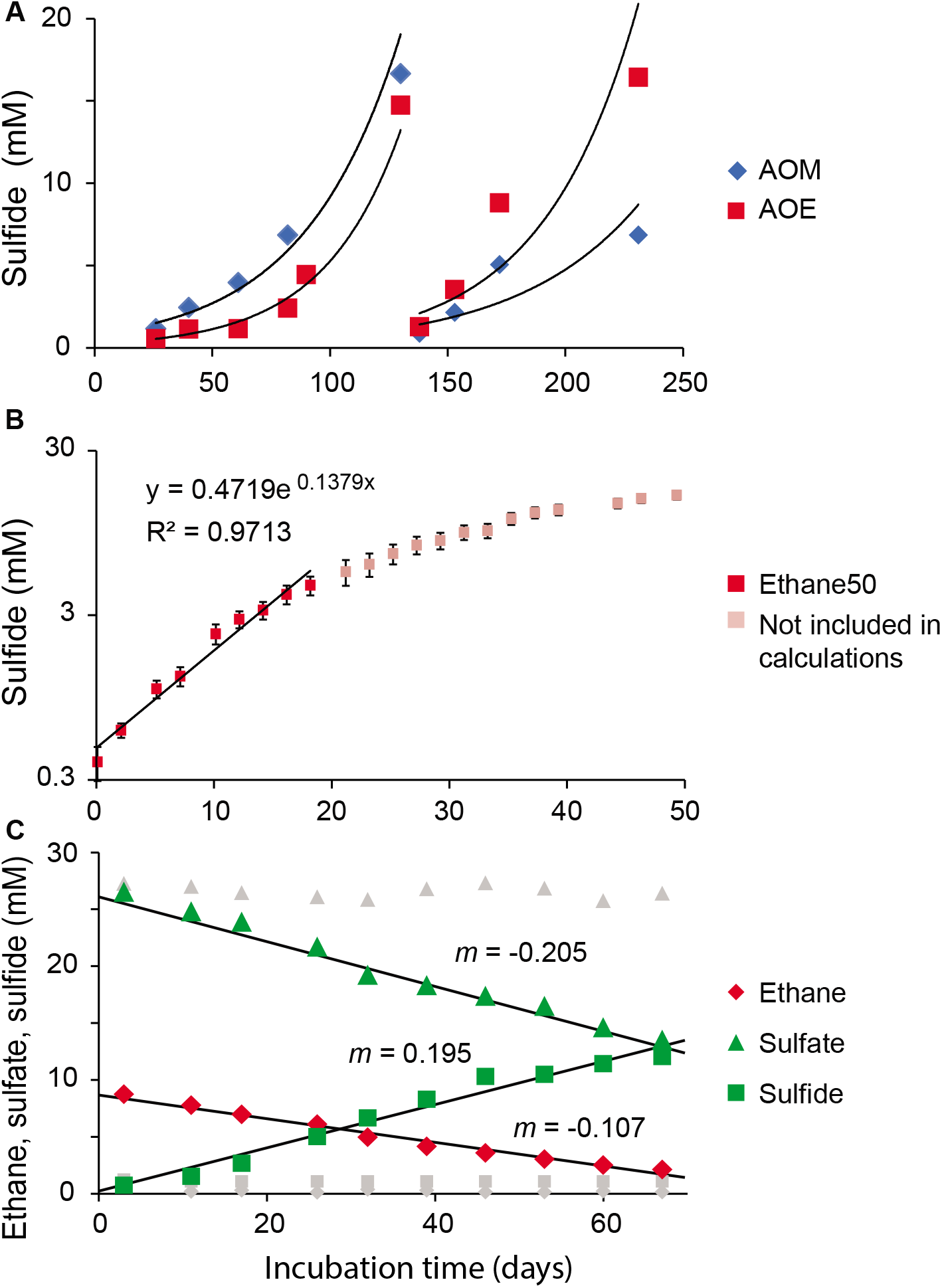
Cultivation and stoichiometry test of the Ethane 50 culture. **A,** Rates of methane-dependent (blue) and ethane-dependent (red) sulfide production in sediments of the Guaymas Basin incubated at 50°C.. **B.** Determination of activity doubling times in anaerobic ethane-oxidizing culture. Logarithmic y-axis with sulfide production shows a decrease in activity at 3 mM sulfide and estimated activity doubling times in low sulfide concentrations of 6-7 days. **C,** Development of ethane (diamond), sulfate (triangles) and sulfide (squares) concentrations in the Ethane50 culture. Gray symbols show corresponding concentrations measured in control incubations without ethane addition (data from 1 of 3 replicate incubations, for complete data see Table S6. The ratios of the slopes of sulfate and sulfide to ethane (1.92 and 1.82) are close to the stoichiometric ratios of sulfate reduction and ethane oxidation. The small offset may relate to biomass production and sampling artifacts.

A stoichiometric growth experiment with the Ethane50 culture (Fig. 1B) showed that ethane is completely oxidized while sulfate was reduced to sulfide according to the formula:

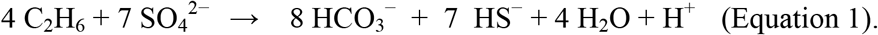

An experiment tracking the exponential development of sulfide over time suggested doubling times of only 6 days at low sulfide concentrations of <5 mM (Fig. 1B), which is substantially faster than estimated for thermophilic AOM consortia with about 60 days [10], and also faster than the cold-adapted anaerobic ethane-oxidizing cultures [5]. Sulfide concentrations over 5 mM seemed to suppress activity and growth of the ethane-oxidizing microorganisms (Fig. 1C). Hence, flow-through bioreactors could be beneficial to increase biomass yields of anaerobic ethane degraders.

### Microbial composition of the Ethane50 culture

Amplified archaeal and bacterial 16S rRNA genes of the original sediment and early, still sediment-containing cultures (150 days of incubation) were sequenced to track the development of microbial compositions over time (for primers, see Table S1). The original sediment contained large number of ANME-1 and the putative partner bacteria *Ca*. Desulfofervidus. The AOM culture got further enriched in ANME-1 archaea and *Ca*. Desulfofervidus, whereas in the Ethane50 culture the GoM-Arc1 clade increased from < 0.1% in the original sediment to roughly 35% of all archaea (Fig. 2A). Notably, the relative abundance of *Ca*. Desulfofervidus increased also in the Ethane50 culture. This indicates that *Ca*. Desulfofervidus was also involved as partner bacterium in the thermophilic ethane culture.

**Fig. 2.**
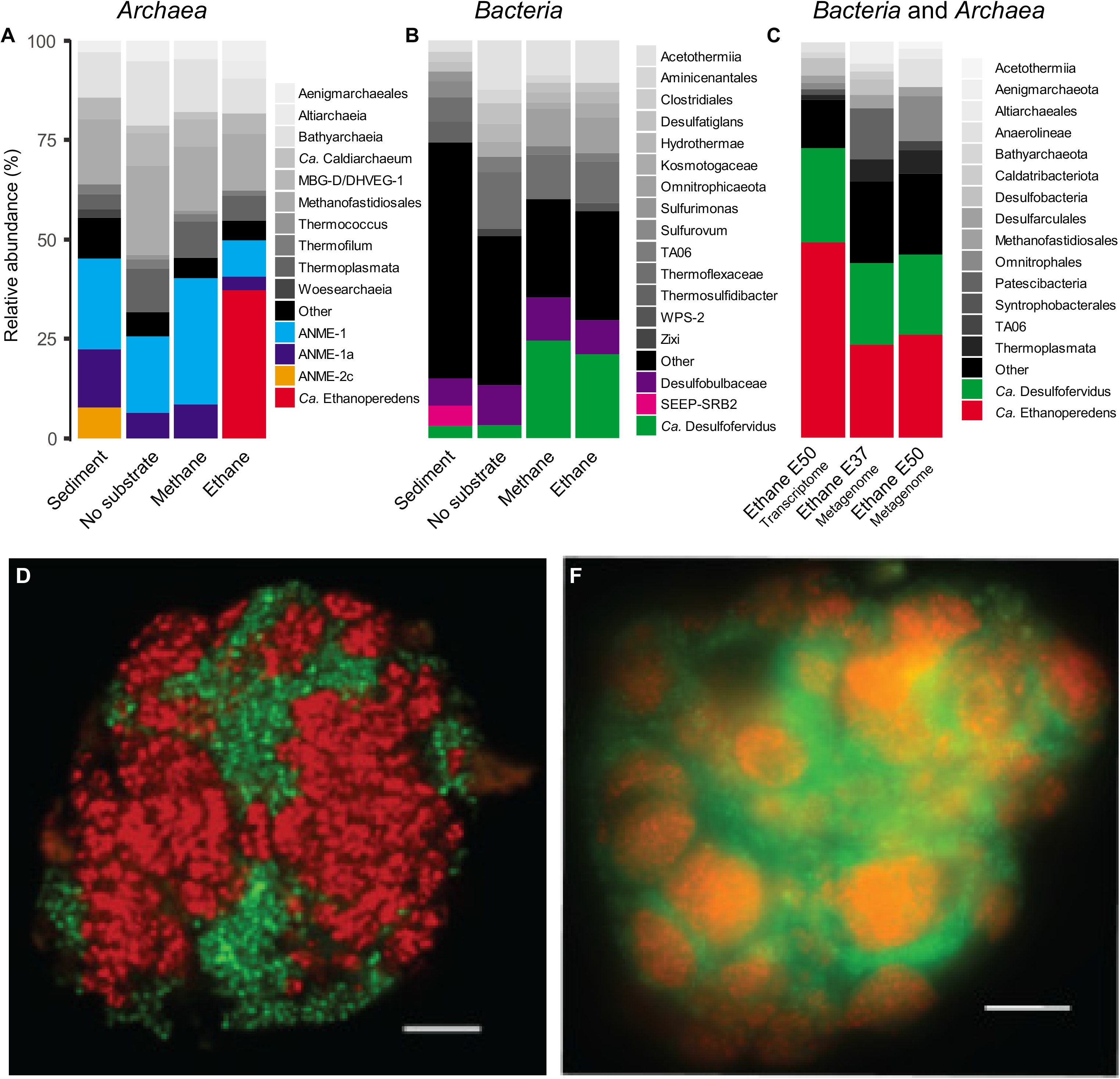
Microbial composition of the Ethane50 culture. **A,** Relative abundance of phylogenetic clades of archaea and **B,** bacteria based on 16S rRNA gene amplicon sequencing present in the inoculated sediment, and in cultures with no substrate, with methane and ethane after 150 days of incubation. **C,** relative abundance of active microbial groups based on 16 rRNA fragments recruited from the genome of Ethane37 and Ethane50 after 2.5 years of incubation and the transcriptome of the Ethane50 culture after one year of incubation with ethane. **D,** Laser-scanning micrograph and **E**, epifluorescence micrograph of microbial consortia stained with probes specific for the GoM-Arc1 clade (red, Alexa-594) and *Ca*. Desulfofervidus (green, Alexa-488) in the Ethane50 culture. Scale 10 μm.

To visualize the cells involved in the anaerobic oxidation of ethane, oligonucleotide probes specific for the GoM-Arc1 clade and *Ca*. Desulfofervidus were applied on the Ethane50 culture using catalyzed reporter deposition fluorescence in situ hybridization (CARD-FISH; for probes see Table S1). The Ethane50 culture contained large and tightly packed consortia with sizes of up to 40 μm in diameter formed by GoM-Arc1 and *Ca*. Desulfofervidus cells (Fig 2D,E). In the consortia, archaea and bacteria grew spatially separated. These large consortia apparently develop from small but already dense consortia found in the inoculate, similar to what was found for cold-adapted AOM consortia [19]. Such a separation of the partner organisms is also characteristic for consortia in the butane-degrading culture [9] and for most AOM consortia [20]. In contrast, in thermophilic AOM consortia of ANME-1 and *Ca*. Desulfofervidus, the partner cells appear well-mixed [21]. The Ethane50 culture differs from the cold-adapted ethane oxidizing culture in which *Ca*. Argoarchaeum forms rather loose assemblages with yet uncharacterized bacteria [5].

To analyze the metabolic potential of the microorganisms involved in ethane degradation, Ethane37 and Ethane50 cultures were subjected to transcriptomic and genomic analysis. The 16S rRNA sequences extracted from the shot-gun RNA reads of the Ethane50 culture were strongly dominated by GoM-Arc1 (60%) and *Ca*. Desulfofervidus (20%; Fig. 2C), supporting a crucial role of these two organisms in thermophilic ethane degradation. Long-read DNA sequencing for the Ethane50 culture resulted in a partial genome of GoM-Arc1 with 76.2% completeness (GoM-Arc1_E50_DN), whereas by applying this approach to the Ethane37 culture we obtained a closed genome of the GoM-Arc1 archaeon (GoM-Arc1_E37). Both GoM-Arc1 genomes share an average nucleotide identity (ANI) of 98%, hence a complete consensus genome for Ethane50 (GoM-Arc1_E50) was obtained by mapping long reads of the Ethane50 culture on the closed GoM-Arc1_E37 genome (see Material and Methods and Table S2). GoM-Arc1_E50 had a size of 1.92 MB and a GC content of 46.5%. To assess the genomic diversity of archaea of the GoM-Arc1 clade, additionally a MAG of GoM-Arc1 from the Loki’s Castle hydrothermal vent field (GoM-Arc1-LC) with a completeness of 68% and eight single-cell amplified genomes (SAGs) form different cold seeps and different completenesses (10 to 59%) were retrieved (Table S2). The MAG GoM-Arc1-LC and the eight single cells have an average nucleotide identity (ANI) of over 90% suggesting that they belong to the same species. The 16S rRNA gene identity is in the range of 99.5% supporting a definition as same species and shows that the same species of GoM-Arc1 can be found in diverse seep sites (Table S2 and Figure S1). Together with several MAGs of the GoM-Arc1 clade archaea from public databases [5, 17, 18] these MAGs now provide an extensive database for the genomic description of the GoM-Arc1 clade. All GoM-Arc1 clade genomes have an estimated size smaller than 2 Mb, which is in the range of the other thermophilic alkane degraders, such as *Ca*. Syntrophoarchaeum (1.5-1.7 Mb) and ANME-1 (1.4-1.8 Mb) [9, 22]. The genome is however much smaller than the 3.5 Mb genome of the mesophilic sister lineage *Ca*. Methanoperedens. This organism is thriving on methane and is capable to reduce nitrate or metals without partner bacteria [23, 24].

All GoM-Arc1 genomes contain the genes encoding the enzymes of the methanogenesis pathway, including a highly similar divergent-type MCR and the Wood-Ljungdahl pathway, but no pathway for beta-oxidation of longer fatty acids. Hence it is likely that all members of this clade are ethane oxidizers. Based on 16S rRNA gene phylogeny and a genome tree based on 32 marker genes, the GoM-Arc1 clade divides into two sub-clusters. According to a 16S rRNA gene identity of ~95% (Fig. S1) and an average amino acid identity (AAI) of ~63% (Fig. 3A; Table S2), these clusters should represent two different genera. One cluster contains the recently described ethane oxidizer *Candidatus* Argoarchaeum ethanivorans and genomes derived from cold environments including the Gulf of Mexico and the moderately heated Loki’s Castle seeps [25]. The second cluster includes the thermophilic GoM-Arc1 strains found in the Ethane50 and the Ethane37 culture and sequences of other MAGs from the Guaymas Basin [16, 18]. Based on the substrate specificity (see results below) and its optimal growth at elevated temperatures, we propose to name the Ethane50 strain of GoM-Arc1 *Candidatus* Ethanoperedens thermophilum (Ethanoperedens, Latin for nourishing on ethane; thermophilum, Latin for heat loving).

**Fig. 3.**
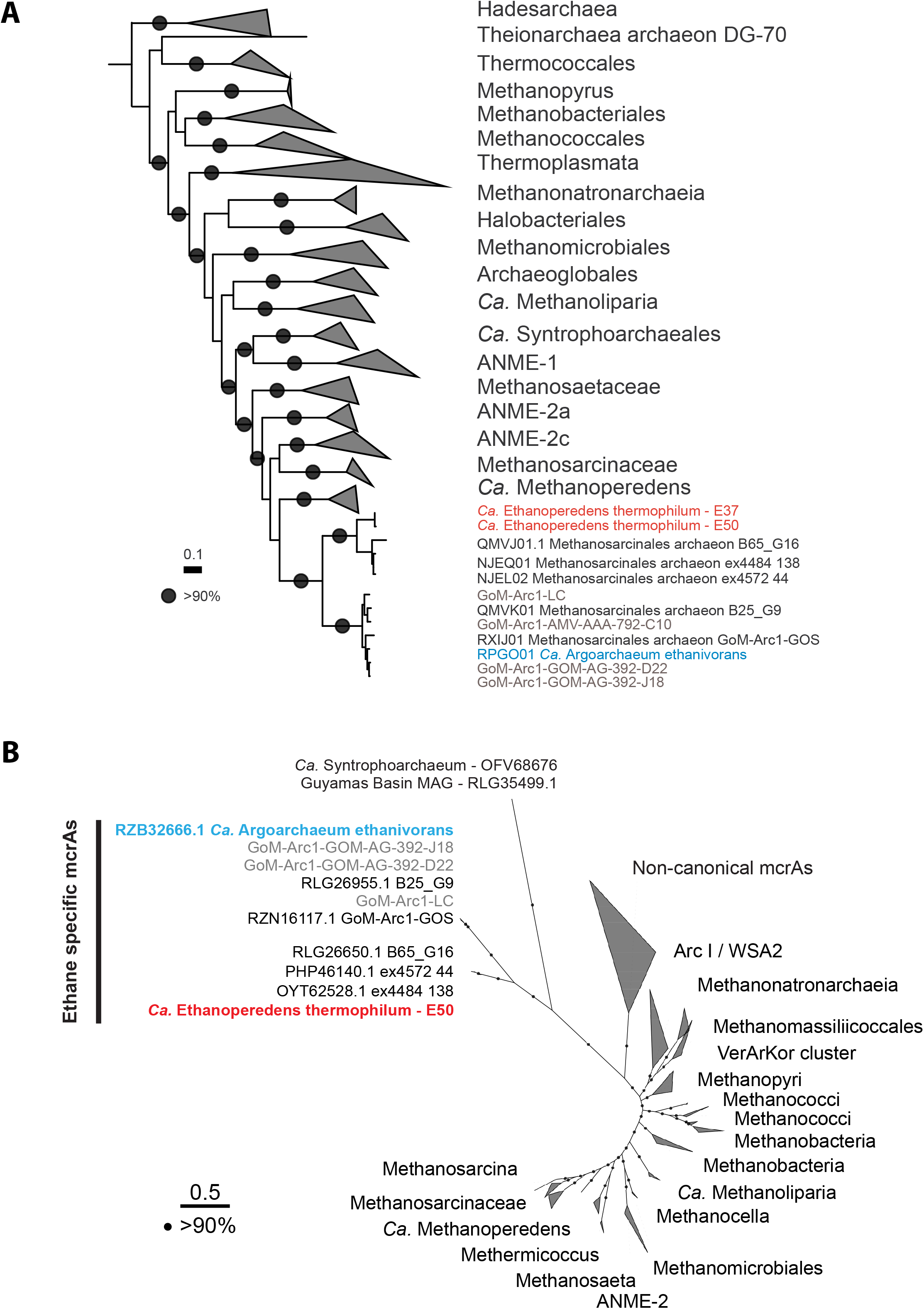
Phylogenetic affiliation based on 32 marker genes and *mcr*A amino acid sequences of *Ca*. Ethanoperedens. **A**, Phylogenetic affiliation of *Ca*. Ethanoperedens within the *Euryarchaeota* based on 32 aligned marker gene amino acid sequences; outgroup is *Thaumarchaeota*. The scale bar indicates 10% sequence divergence. **B**, Phylogenetic affiliation of *mcr*A amino acid sequences. The *mcr*A sequences of GoM-Arc1 form a distinct branch within the non-canonical, potentially multi-carbon alkane-activating MCRs. The *mcr*A genes of the GoM-Arc1 cluster can be further divided into those from cold-adapted organisms, including *Ca*. Argoarchaeum ethanivorans, and the cluster including the thermophiles of the genus *Ca*. Ethanoperedens. Sequences from the Ethane50 enrichment are depicted in red, environmental sequences from metagenomes and single cell genomes from this study in grey and *Ca*. Argoarchaeum ethanivorans in blue.

### Genomic and catabolic features of *Ca*. Ethanoperedens

The main catabolic pathways of *Ca*. Ethanoperedens are a complete methanogenesis and a Wood-Ljungdahl pathway (Fig. 4). Its genome encodes for only one MCR. The three MCR subunits αβγ are on a single operon. The amino acid sequence of the single alpha subunit (*mcr*A) of *Ca*. Ethanoperedens is phylogenetically most closely related with the recently described divergent-type MCR of *Ca*. Argoarchaeum with an amino acid identity of 69%, but also with all other *mcr*A sequences of GoM-Arc1 archaea [5, 12, 16, 18]. These MCRs form a distinct cluster in comparison to other divergent MCRs and to the canonical MCRs of methanogens and methanotrophs (Fig. 3B).

**Fig. 4.**
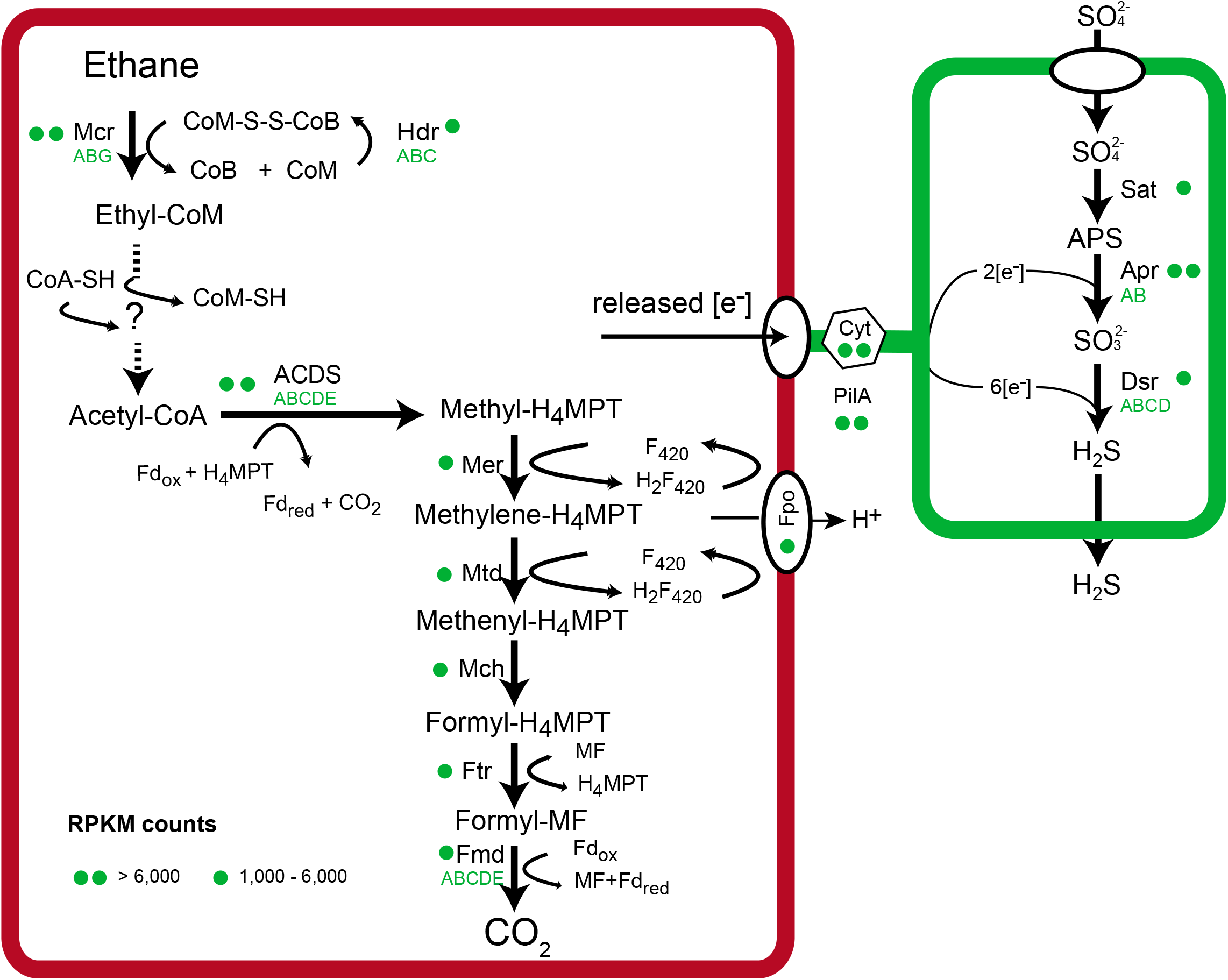
Metabolic model of anaerobic ethane oxidation in *Ca*. Ethanoperedens thermophilum. Ethane is activated in the ethane specific MCR. The produced CoM-bound ethyl groups are consecutively oxidized and transformed to CoA-bound acetyl units. Acetyl-CoA is cleaved using the ACDS of the Wood-Ljungdahl pathway. The remaining methyl groups are fully oxidized on the reversed methanogenesis pathway. Similar to ANME archaea and *Ca*. Syntrophoarchaeum, *Ca*. Ethanoperedens does not contain a reductive pathway; hence electrons released during ethane oxidation are transferred to the partner bacterium *Ca*. D. auxilii. Therefore, in both partners, cytochromes and pili are present and expressed, similar as described in thermophilic consortia performing AOM [22] (for detailed expression patterns, see Table S3)

The similarity of GoM-Arc1 *mcr*A sequences to the described canonical and non-canonical sequences is below 43% and changes in the amino acid sequences are also found in the highly conserved active site of the enzyme (Fig. S2). The relative expression of the *mcr* subunits compared to all reads mapping to *Ca*. Ethanoperedens (reads per kilo base per million mapped reads; RPKM; i.e. *mcr*A = 9790) is at least two times higher than the expression of all other genes of the main catabolic pathway (Fig. 4; Table S3). The relative *mcr* expression of *Ca*. Ethanoperedens is higher than the expression of the multiple *mcr* genes in *Ca*. Syntrophoarchaeum, but lower than the expression of *mcr* in thermophilic ANME-1 archaea [9, 22]. The relatively low expression of *mcr* in short-chain alkane oxidizing archaea can be explained by the properties of their substrates. Short-chain alkane oxidation releases larger amounts of energy than methane oxidation. Furthermore, the cleavage of C-H bonds in multi-carbon compounds requires less energy than the cleavage of C-H bonds of methane [26], hence less MCR might be required to supply the organism with sufficient energy.

To test the substrates activated by the MCR of *Ca*. Ethanoperedens, we supplied different alkanes to the active Ethane50 culture replicates and analysed the extracted metabolites. Cultures supplied with ethane show the *m/z* 168.9988 of the authentic ethyl-CoM standard (Fig. 5A,B), which was not observed in the control incubation without substrate. Moreover, addition of 30% 1-^13^C-ethane resulted in the increase of masses expected for 1-^13^C-ethyl-CoM and 2-^13^C-ethyl-CoM (Fig. 5C). This confirms that *Ca*. Ethanoperedens produces ethyl-CoM from ethane. To test substrate specificity of *Ca*. Ethanoperedens, we provided culture replicates with four different gaseous alkanes (methane, ethane, propane and *n*-butane, and a mix of all four substrates). Besides the ethane-amended culture, sulfide was only produced in the Ethane50 culture supplied with the substrate mix (Fig. S3). In agreement to this, no other alkyl-CoM variant apart from ethyl-CoM was detected (Fig. 5A). This shows that the MCR of *Ca*. Ethanoperedens and most likely all MCR enzymes of GoM-Arc1 archaea (Fig. 3B) activate ethane, but no or only trace amounts of methane and other alkanes. The high substrate specificity of the MCR is crucial for GoM-Arc1 archaea, as they miss the enzymatic machinery to use larger CoM-bound intermediates, since they lack the fatty acid degradation pathway that is required to degrade butane and propane [9]. *Ca*. Ethanoperedens contains and expresses a complete methyl transferase (*mtr*). This enzyme might cleave small amounts of methyl-CoM that might be formed as side reaction of the MCR. The methyl unit would be directly transferred to the methylene-H4MPT reductase (*mer*) and oxidized in the upstream part of the methanogenesis pathway to CO_2_ (Fig. 4).

**Fig. 5.**
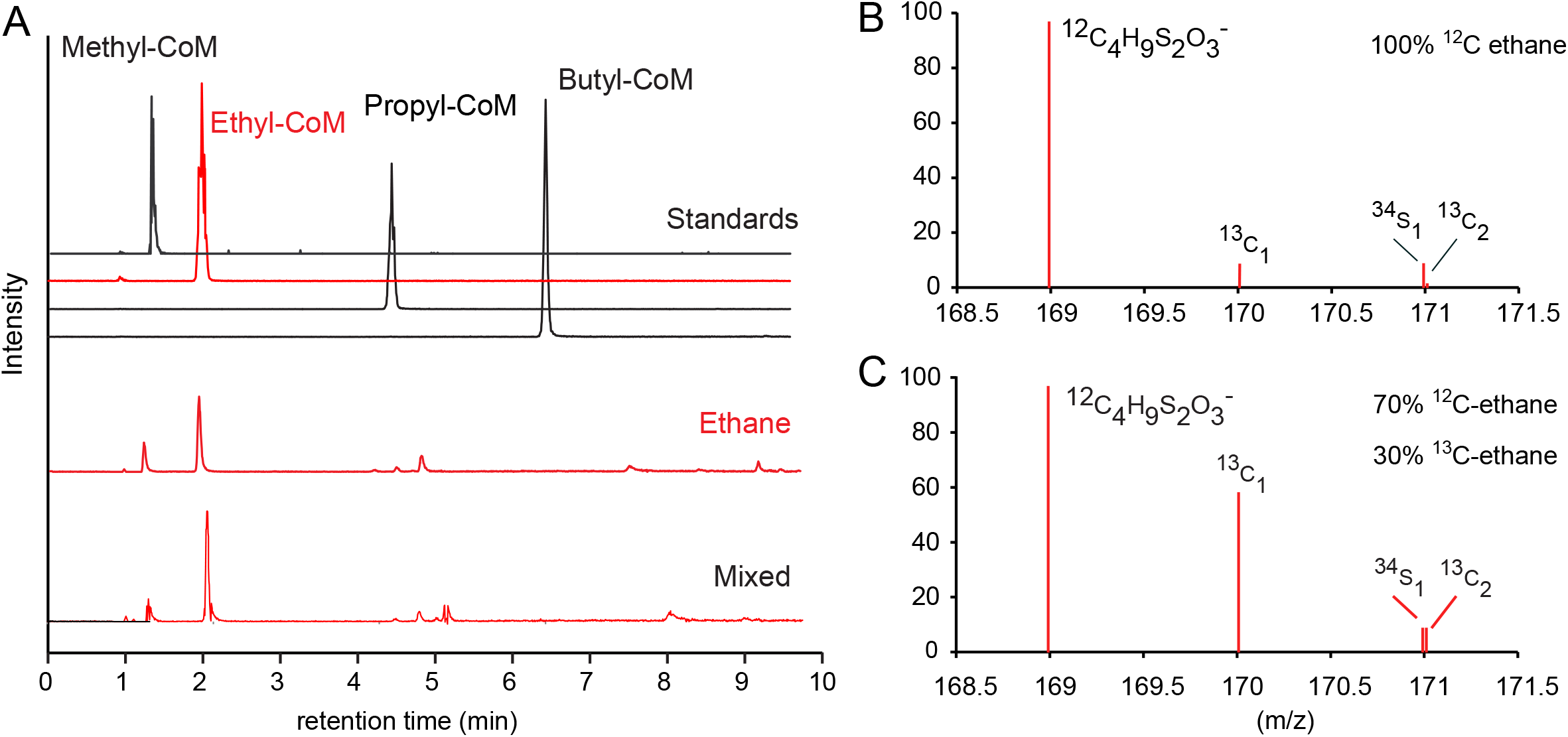
Detection of coenzyme M bound intermediates in the Ethane50 culture. **A.** Upper 4 panels, Total ion counts for HPLC peaks for authentic standards of methyl-, ethyl-, propyl and butyl CoM, and chromatograms for ethane and mixed alkane gases (methane to butane). **B,C.** Mass spectra (*m/z* 168.5-171.5) for culture extracts after providing the Ethane50 culture with **(B)** non-labeled ethane and **(C)** 30% ^13^C-labeled ethane. Diagrams show the relative intensities for ethyl-CoM-H (^12^C_4_H_9_S_2_O_3_^-^) calculate *m/z* 168.9988) and its isotopomers with one or two ^13^C carbon or one ^34^S isotope.

Based on the observed net reaction and the genomic information, *Ca*. Ethanoperedens completely oxidizes ethane to CO_2_. In this pathway, coenzyme A bound acetyl units are oxidized in the Wood-Ljungdahl pathway and the upstream part of the methanogenesis pathway (Fig. 4). Our model, however, does not explain how CoM-bound ethyl groups are oxidized to acetyl units and ligated to CoA. Similar transformations are required in the other multi-carbon alkane-oxidizing archaea such as *Ca*. Syntrophoarchaeum and *Ca*. Argoarchaeum [5, 9]. Those oxidation reactions lack biochemical analogues, hence genomic information alone allows only indirect hints on their function. In *Ca*. Ethanoperedens, a release of ethyl-units and transformation as free molecules (ethanol to acetate) is unlikely, because a subsequently required formation of acetyl-CoA from acetate would require CoA ligases, which are not present in the genome. Instead, the transformation of ethyl into acetyl units could be performed by a tungstate-containing aldehyde ferredoxin oxidoreductase (AOR) that could catalyse the oxidation with cofactors such as CoM or CoA. In the archaeon *Pyrococcus furiosus* AORs transform aldehydes to the corresponding carboxylic acid [27]. Both, *Ca*. Ethanoperedens and *Ca*. Argoarchaeum genomes, contain three *aor* copies, and in all cases these genes are either located in close proximity or on operons with genes of the methanogenesis pathway. We detected a high expression of two of the three *aor* genes (RPKM *aor* = 3805 and 7928), indicating a viable function of the enzymes. Likewise, very high protein concentrations of these enzymes were shown for *Ca*. Argoarchaeum [5], supporting the hypothesis of a critical function. An *aor* gene is also present in the butane oxidizer *Ca*. Syntrophoarchaeum, yet its expression is rather moderate [9], which questions its role in the catabolic pathway of this organism. In contrast ANME archaea do not contain or overexpress *aor* genes, likely because the encoded enzymes have no central role in their metabolism. We searched the cell extracts for potential intermediates in the pathway, but based on retention time and mass we were not able to detect potential intermediates such as ethyl-CoA. Similarly, acetyl-CoA, the substrate of the Wood-Ljungdahl pathway, was not detected. A lack of detection however does not exclude those compounds as intermediate. Instead, the compound turnover might be very fast, which could be required for an efficient net reaction. Additionally, a mass spectrometric detection of unknown intermediates could be hindered by compound instability or loss during the extraction. Further metabolite studies and enzyme characterizations are required to understand the role of AOR in alkane oxidation

Acetyl-CoA, the product formed of the above proposed reactions, can be introduced into the Wood-Ljungdahl pathway. The acetyl group is decarboxylated by the highly expressed acetyl-CoA decarbonylase/synthase (ACDS), and the remaining methyl group is transferred to tetrahydromethanopterin (H_4_-MPT). The formed methyl-H_4_-MPT can then be further oxidized to CO_2_ following the reverse methanogenesis pathway (Fig. 4). *Ca*. Ethanoperedens lacks genes for sulfate or nitrate reduction, similarly to other genomes of the GoM-Arc1 clade. The electrons produced in the oxidation of ethane thus need to be transferred to the sulfate-reducing partner bacterium *Ca*. D. auxilii, as previously shown for the anaerobic oxidation of methane and butane. In co-cultures of *Ca*. Argoarchaeum and their partner bacteria, Chen and co-workers (2019) suggest the transfer of reducing equivalents via zero-valent sulfur between the loosely aggregated *Ca*. Argoarchaeum and its partner bacterium, analogous to the hypothesis of Milucka, Ferdelman [28]. In the Ethane50 culture such a mode of interaction is highly unlikely, as the partner *Ca*. D. auxilii is an obligate sulfate reducer, incapable of sulfur disproportionation [11]. Based on genomic information, direct electron transfer appears to be more likely. Alkane-oxidizing archaea and their partner bacterium *Ca*. D. auxilii produce cytochromes and pili-based nanowires when supplied with their substrate [9, 29, 30]. Also *Ca*. Ethanoperedens contains 11 different genes for cytochromes with expression values of up to 14800 RPKM representing some of the highest expressed genes in the culture (Table S3). Interestingly, *Ca*. Ethanoperedens also contains and expresses a type IV pilin protein with a high RPKM value of 11246. The partner bacterium *Ca*. Desulfofervidus also shows a high expression of pili and cytochromes under ethane supply, showing their potential importance for the interaction of these two organisms in the syntrophic coupling of ethane oxidation to sulfate reduction.

### Environmental distribution of GoM-Arc1 archaea

16S rRNA gene sequences clustering with *Ca*. Ethanoperedens and *Ca*. Argoarchaeum have been found in hydrocarbon-rich marine environments like cold seep and hot vent environments, including asphalt seeps in the Gulf of Mexico and the Guaymas Basin hydrothermal vents in the Gulf of California [31–33]. In some environments like oil seeps of the Gulf of Mexico and gas-rich barite chimneys of the Loki’s Castle, 16S rRNA gene surveys have shown that up to 30% of archaeal gene sequences belonged to the GoM-Arc1 clade [12]. To estimate absolute abundances and potential partnerships of GoM-Arc1 in the environment, we performed CARD-FISH on samples from different seep and vent sites across the globe (Fig. 6). With up to 10^8^ cells per ml, archaea of the GoM-Arc1 clade were particularly abundant in cold seep sediments in the northern Gulf of Mexico (St. 156). This cold seep transports thermogenic hydrocarbon gases that are particularly enriched in short-chain alkanes [34, 35]. Other cold seep and hot vent sediments from the Guaymas Basin, Hydrate Ridge and Amon Mud Volcano contain between 10^5^ and 10^6^ GoM-Arc1 cells per ml of sediment, which represents 1-5% of the archaeal community (Fig. 6A). At all sites we found that GoM-Arc1 associates with partner bacteria. At the hydrothermally-heated site in the Guaymas Basin, GoM-Arc1 aggregated with *Ca*. Desulfofervidus, the partner bacterium of the Ethane37 and Ethane50 cultures. At Loki’s Castle, GoM-Arc1 and *Ca*. Desulfofervidus were co-occurring in barite chimneys based on sequence information, yet they were not found to form the same tight consortia as at other sites. At the temperate site Katakolo Bay in Greece, GoM-Arc1 archaea formed consortia with very large, yet unidentified vibrioform bacteria (Fig. 6B-F). These cells hybridized with a probe for *Deltaproteobacteria* but not with probes for known partner bacteria (for probes see Table S1). At the cold seep sites, the associated cells could not be stained with probes for the known partner bacteria of cold-adapted ANME including SEEP-SRB1, and SEEP-SRB2, and also not with that for *Ca*. Desulfofervidus. It remains an important question as to how the archaea can select only few specific types of bacteria as partner in the anaerobic alkane oxidation, and for which specific traits they are selected. Based on their global presence in hydrocarbon-rich environments, GoM-Arc1 archaea could be considered as key player in the anaerobic oxidation of ethane in marine sediments. Its role would be similar to the role of ANME archaea in AOM.

**Fig. 6.**
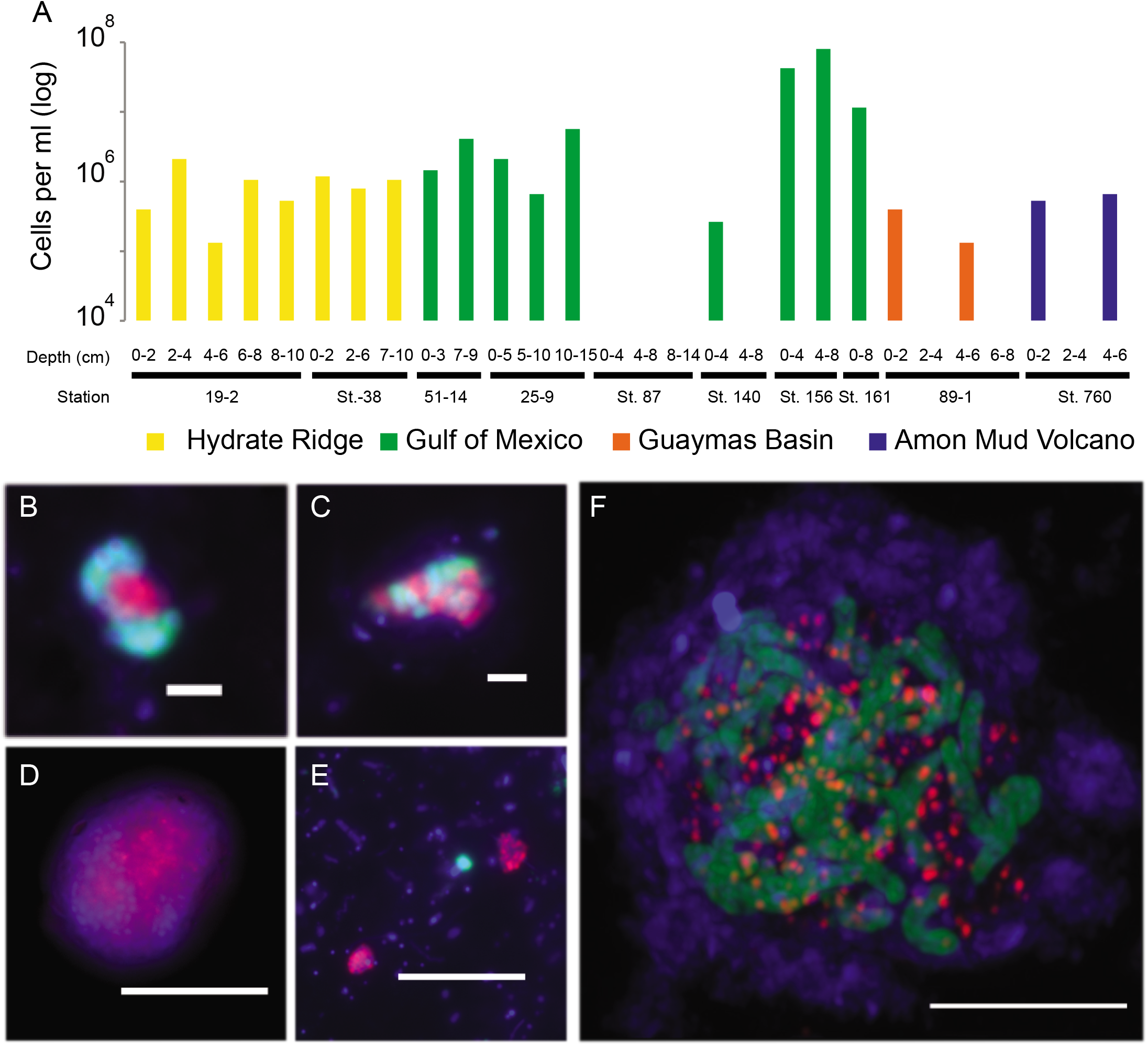
Abundance and exemplary micrographs of GoM-Arc1 archaea in sediments from cold seeps and Guaymas Basin. **A**. Abundance estimations of archaeal cells detected by the GoM-Arc1 specific probe GOM-ARCI-660 in a CARD-FISH survey. Detection limit approx. 5×10^4^ cells per ml sediment. **B – F** Epifluorescence (**B-E**) and laser-scanning micrographs (**F**) of environmental samples using CARD-FISH with combining the GoM-Arc1 specific probe (red) and the general bacterial probe EUB-338 (green). Environmental samples originated from the seep sites Hydrate Ridge, Oregon (**B**); Gulf of Mexico (**C**), Guaymas Basin (**D**); Loki’s Castle (**E**) and Katakolo Bay, Greece (**F**). Scale 5 μm for D-F and 2 μm for B and C.

### Future possible applications of *Ca*. Ethanoperedens

Archaea of the GoM-Arc1 cluster are likely the dominant, if not the only organisms capable of the anaerobic oxidation of ethane in the global seafloor. An important further task is to assess deep oil and gas reservoirs for their diversity of ethane oxidizers. The rapid growth of *Ca*. Ethanoperedens and the streamlined genome make it a model organism for the study of anaerobic ethanotrophy in archaea. The biochemistry of short-chain alkane-oxidizing archaea will be of high interest for future biotechnological applications. An organism using the metabolism of *Ca*. Ethanoperedens in the reverse direction should be able to produce ethane, similar to methane production by methanogens. Yet, there is scarce isotopic evidence for the existence of ethanogenic organism in nature [36]. Furthermore, under common environmental conditions thermodynamics favor the production of methane from inorganic carbon over the production of ethane. To test the general reversibility of the ethane oxidation pathway, we incubated the active Ethane50 culture with ^13^C-labelled inorganic carbon and traced the label transfer into ethane. Within 18 days δ^13^C-ethane values increased from −3‰ to +120‰, whereas isotopic compositions in the non-labeled culture remained stable (Fig. S4). Considering the forward rate and ethane stock, the back reaction amounts to 1.5‰ to 3% of the forward reaction, which is in the range for back fluxes of carbon measured in AOM [21, 37]. This experiment shows that the ethane oxidation pathway is fully reversible. To test the net ethane formation in the Ethane50 culture, we removed sulfate from culture aliquots and added hydrogen as electron donor. These cultures formed between 1 and 17 μmol 1^−1^ ethane within 27 days (Table S6). The ethane production was however a very small fraction of 0.08% of the ethane oxidation rate in replicate incubations with ethane and sulfate. No ethane was formed in the presence of hydrogen and sulfate. We interpret the ethane formation in the culture as enzymatic effect in the ethane-oxidizing consortia. Hydrogenases will fuel reducing equivalents into the pathway, which may ultimately lead to the reduction of carbon dioxide to ethane. A growing culture could not be established under these conditions, however, the experiments suggest that related or genetically modified methanogenic archaea could thrive as ethanogens. A complete understanding of the pathway and enzymes of GoM-Arc1 archaea, however, is required to develop the biotechnological potential of an ethanogenic organism. To allow energy-conserving electron flows in this organism, a genetically modified methanogen should be used as host organism. For a targeted modification of such archaea, the pathway of ethane oxidation must be completely understood, and research should focus especially on the transformation of coenzyme M bound ethyl units to coenzyme A bound acetyl units.

## MATERIAL AND METHODS

### Inoculum and establishment of alkane-oxidizing cultures

This study bases on samples collected during R/V ATLANTIS cruise AT37-06 with submersible *Alvin* to the Guaymas Basin vent area in December 2016 (for locations see Table S4). A sediment sample was collected by push coring within a hydrothermal area marked by conspicuous orange-type *Beggiatoa* mats (Dive 4869, Core 26, 27°0.4505’N 111°24.5389’W, 2001 m water depth, December 20, 2016). The sampling site was located in the hydrothermal area where, during a previous *Alvin* visit, sediment cores containing locally ^13^C-enriched ethane had indicated ethane-oxidizing microbial activity [33]. In situ temperature measurements using the *Alvin* heat flow probe revealed a steep temperature gradient reaching 80°C at 30-40 cm sediment depth. The retrieved samples contained large amounts of natural gas as observed by bubble formation. Soon after recovery, the overlying *Beggiatoa* mat was removed, and the top 10 cm of the sediment were filled into 250 ml Duran bottles, which were gas-tight sealed with butyl rubber stoppers. In the home laboratory, sediments were transferred into an anoxic chamber. There a sediment slurry (20% sediment and 80% medium) was produced with synthetic sulfate reducer (SR) medium (pH 7.0) [38, 39], and distributed into replicate bottles (sediment dry weight per bottle 1.45 g). These bottles were amended with methane or ethane (0.2 MPa), or kept with a N_2_ atmosphere without alkane substrate. These samples were incubated at 37°C, 50°C and 70°C. To determine substrate-dependent sulfide production rates, sulfide concentrations were measured every 2-4 weeks using a copper sulfate assay [40]. Ethane-dependent sulfide production was observed at 37°C and 50°C, but not at 70°C. When the sulfide concentration exceeded 15 mM the cultures were diluted (1:3) in SR medium and resupplied with ethane. Repeated dilutions led to virtually sediment-free, highly active cultures within 18 months. A slight decrease of the initial pH value to 6.5 led to increased ethane oxidation activity and faster growth in the culture.

### Quantitative substrate turnover experiment

The Ethane50 culture was equally distributed in 6 × 150 ml serum flasks using 20 ml inoculum and 80 ml medium. Three replicate cultures were amended with 0.05 MPa ethane in 0.1 MPa N_2_:CO_2_, while 3 negative controls were amended with 0.15 MPa N_2_:CO_2_. Both treatments were incubated at 50°C. Weekly, 0.5 ml headspace gas samples were analyzed for ethane content using an Agilent 6890 gas chromatograph in split-less mode equipped with a packed column (Supelco Porapak Q, 6ft×1/8’× 2.1 mm SS, oven temp 80°C). The carrier gas was helium (20 ml per minute) and hydrocarbons were detected by flame ionization detection. Each sample was analyzed in triplicates and quantified against ethane standards of 5, 10 and 100%. Derived concentrations were converted into molar amounts by taking the headspace size, pressure and temperature into account. Results were corrected for sampled volumes. Sulfide concentrations were measured as described above. To determine sulfate concentrations 1 ml of sample was fixed in 0.5 ml zinc acetate. Samples were diluted 1:50 with deionized water (MilliQ grade; >18.5 MΩ) and samples were measured using non-suppressed ion chromatography (Metrohm 930 Compact IC Metrosep A PCC HC/4.0 preconcentration and Metrosep A Suup 5 −150/4.0 chromatography column).

### DNA extraction, 16S rRNA gene amplification and tag sequencing

DNA was extracted from the different cultures and the original sediment with the MoBio power soil DNA extraction kit (MO BIO Laboratories Inc., Carlsbad, USA) using a modified protocol. 20 ml of the culture was pelleted via centrifugation (5.000 × g; 10 min). The pellet was resuspended in saline phosphate buffer (PBS) and transferred to the *PowerBeat Tube* (manufacturer information needed). The cells were lysed by three cycles of freezing in liquid nitrogen (20 sec) and thawing (5 min at 60°C). After cooling down to room temperature, 10 μl of proteinase K (20 mg ml^−1^) were added and incubated for 30 min at 55°C. Subsequently, 60 μl of solution C1 (contains SDS) were added and the tubes were briefly centrifuged. The samples were homogenized 2 times for 30 sec at 6.0 M/S using a FastPrep-24 instrument (MP Biomedicals, Eschwege, Germany). In between the runs, the samples were kept on ice for 5 min. After these steps, the protocol was followed further according to the manufacturer’s recommendations. DNA concentrations were measured using a Qubit 2.0 instrument (Invitrogen, Carlsbad, USA). 2 ng of DNA was used for amplicon PCR and the product used for 16S rDNA amplicon library preparation following the 16S Metagenomic Sequencing Library Preparation guide provided by Illumina. The Arch349F - Arch915R primer pair was used to amplify the archaeal V3-V5 region and the Bact341F - Bact785R primer pair for the bacterial V3-V4 region (Table S1). Amplicon libraries for both Archaea and Bacteria were sequenced on an Illumina MiSeq instrument (2×300 bp paired end run, v3 chemistry) at CeBiTec (Bielefeld, Germany). After analysis adapters and primer sequences were clipped from the retrieved sequences using cutadapt [41](v1.16) with 0.16 (-e) as maximum allowed error rate and no indels allowed. Resulting reads were analysed using the SILVAngs pipeline using the default parameters (https://ngs.arb-silva.de/silvangs/) [42–44].

### Extraction of high quality DNA, library preparation and sequencing of genomic DNA

Biomass from 200 ml of the Ethane50 and Ethane37 cultures was pelleted by centrifugation and resuspended in 450 μl of extraction buffer. Genomic DNA was retrieved based on a modified version of the protocol described in [45], including three extraction steps. Resuspended pellet was frozen in liquid N_2_ and thawed in a water bath at 65°C Another 1350 μl of extraction buffer were added. Cells were digested enzymatically by proteinase K (addition of 60 μl 20 mg/ml, incubation at 37°C for 1.5 h under constant shaking at 225 rpm), and chemically lysed (addition of 300 μl 20% SDS for 2 h at 65°C). Samples were centrifuged (20 min, 13.000 × g) and the clear supernatant transferred to a new tube. 2 ml of chloroform: isoamylalcohol (16:1, v/v) were added to the extract, mixed by inverting and centrifuged for 20 minutes at 13.000 × g. The aqueous phase was transferred to a new tube, mixed with 0.6 volumes of isopropanol, and stored over night at −20 ° C for DNA precipitation. The DNA was redissolved in water at 65°C for 5 min and then centrifuged for 40 min at 13.000 × g. The supernatant was removed and the pellet washed with ice-cold ethanol (80%) and centrifugation for 10 min at 13.000 x g. The ethanol was removed and the dried pellet was resuspended in PCR-grade water. This procedure yielded 114 μg and 145 μg high quality genomic DNA from the Ethane37 and the Ethane50 culture, respectively. Samples were sequenced with Pacific Biosciences Sequel as long amplicon (4 – 10 kb) and long read gDNA library at the Max Planck-Genome-Centre (Cologne, Germany). To evaluate the microbial community we extracted 16S rRNA gene reads using Metaxa2 [46] and taxonomically classified them using the SILVA ACT online service [47]. For assembly either HGAP4 (Implemented in the SMRTlink software by PacBio) or Canu (https://github.com/marbl/canu) were used. The closed GoM-Arc1 genome from the Ethane37 culture was prepared manually by the combination of assemblies from the two before mentioned tools. The final genome was polished using the resequencing tool included in the SMRT Link software by PacBio. For not circularized de-novo genomes, the resulting contigs were mapped via minimap2 (https://github.com/lh3/minimap2; parameter: ‘-x asm10’) to a reference genome. The reference consensus genomes were prepared using the resequencing tool implemented in the SMRTLink software of PacBio using either the circular GoM-Arc1 de-novo genome from this study or the publically available *Ca*. Desulfofervidus genome (Accession: NZ_CP013015.1) as reference. Final genomes were automatically annotated using Prokka [48] and the annotation refined manually using the NCBI Blast interface [49]. Average nucleotide and amino acid identities were calculated using Enveomics tools [50].

### Single cell genomics

Anoxic sediment aliquots were shipped to the Bigelow Laboratory Single Cell Genomics Center (SCGC; https://scgc.bigelow.org). Cells were separated, sorted and lysed, and total DNA was amplified by multiple displacement amplification. Single cell DNA was characterized by 16S rRNA gene tag sequences [12, 51]. The single cell amplified DNA from Gulf of Mexico was analyzed sequenced as described before in [12]. Single cell amplified DNA from Amon Mud Volcano AAA-792_C10 was sequenced with Hiseq3000 and MiSeq technology and reads were assembled using SPAdes [52] with the single cell mode. Assembled reads were binned based on tetra-nucleotides, coverage and taxonomy using MetaWatt [53]. The final SAG was evaluated for completeness and contamination using CheckM [54]. Genome annotation was performed as described above.

### Extraction of RNA, reverse transcription, sequencing and read processing

Extraction and sequencing of total RNA was prepared in triplicates. RNA was extracted from 150 ml active Ethane50 culture grown in separate bottles at 50°C. Total RNA was extracted and purified as described in [9] using the Quick-RNA MiniPrep Kit (Zymoresearch, Irvine, CA, USA) and RNeasy MinElute Cleanup Kit (QIAGEN, Hilden, Germany). Per sample at least 150 ng of high-quality RNA were obtained. RNA library was prepared with the TrueSeq Stranded Total RNA Kit (Illumina). An rRNA depletion step was omitted. The samples were sequenced on an Illumina Nextseq with v2 chemistry and 1×150bp read length. The sequencing produced ~50 Gb reads per sample. Adaptors and contaminant sequences were removed and reads were quality trimmed to Q10 using bbduk v36.49 from the BBMAP package. For phylogenetic analysis of the active community 16S rRNA reads were recruited and classified based on SSU SILVA Release 132 [47] using phyloFlash [55]. Trimmed reads were mapped to the closed genomes of the *Candidatus* Ethanoperedens thermophilum and *Ca*. Desulfofervidus using Geneious Prime 2019.2.1 (https://www.geneious.com) with a minimum mapping quality of 30%. The expression level of each gene was quantified by counting the number of unambiguously mapped reads per gene using Geneious. To consider gene length, read counts were converted to reads per kilobase per million mapped reads (RPKM).

### Phylogenetic analysis of 16S rRNA genes, marker genes and *mcr*A amino acid sequences

A 16S rRNA gene based phylogenetic tree was calculated using publically available 16S rRNA sequences from the SSU Ref NR 128 SILVA database [42]. The tree was constructed using ARB [56] and the FastTree 2 package [57] using a 50% similarity filter. Sequence length for all 16S rRNA genes was at least 1100 bp. After tree calculation, partial sequences retrieved from single cells were included into the tree. ARB [56] was used for visualization of the final tree. The marker gene tree was calculated using 126 publically available genomes and genomes presented in this study. The tree was calculated based on aligned amino acid sequences of 32 marker genes picked from known archaeal marker genes (Table S5) [58]. For the preparation of the aligned marker gene amino acid sequences we used the phylogenomic workflow of Anvio 5.5 [59]. The marker gene phylogeny was calculated using RaxML version 8.2.10 [60] with the PROTGAMMAAUTO model and LG likelihood amino acid substitution. 1000 fast bootstraps were calculated to find the optimal tree according to RaxML convergence criterions. The software iTol v3 was used for tree visualization [61]. The *mcr*A amino acid phylogenetic tree was calculated using 358 sequences that are publically available or presented in this study. The sequences were manually aligned using the Geneious Prime 2019.2.1 (https://www.geneious.com) interface and 1060 amino acid positions considered. The aligned sequences were masked using Zorro (https://sourceforge.net/projects/probmask/) and a phylogenetic tree was calculated using RaxML version 8.2.10 [60] using the PROTGAMMAAUTO model and LG likelihood amino acid substitution. 1000 fast bootstraps were calculated. The tree was visualized with iTol v3 [61].

### Catalyzed reported deposition fluorescence in situ hybridization (CARD-FISH)

Aliquots of the Ethane50 culture and environmental samples were fixed for 1 h in 2% formaldehyde, washed three times in PBS (pH = 7.4): ethanol 1:1 and stored in this solution. Aliquots were sonicated (30 sec; 20% power; 20% cycle; Sonoplus HD70; Bandelin) and filtered on GTTP polycarbonate filters (0.2 μm pore size; Millipore, Darmstadt, Germany). CARD-FISH was performed according to [62] including the following modifications: Cells were permeabilized with a lysozyme solution (PBS; pH 7.4, 0.005 M EDTA pH 8.0, 0, 02 M Tris-HCl pH 8.0, 10 mg ml^−1^ lysozyme; Sigma-Aldrich) at 37°C for 60 minutes followed by proteinase K solution treatment (7.5 μg ml^−1^ proteinase K; Merck, Darmstadt, Germany in PBS; pH=7.4, 0.005 M EDTA pH 8.0,0.02 M Tris-HCl pH 8.0,) at room temperature for 5 minutes. Endogenous peroxidases were inactivated by incubation in a solution of 0.15% H_2_O_2_ in methanol for 30 min at room temperature. Horseradish-peroxidase (HRP)-labeled probes were purchased from Biomers.net (Ulm, Germany). Tyramides were labeled with Alexa Fluor 594 or Alexa Fluor 488. All probes were applied as listed in Table S1. For double hybridization, the peroxidases from the first hybridization were inactivated in 0.15% H2O2 in methanol for 30 min at room temperature. Finally, the filters were counterstained with DAPI (4’, 6’ -diamino-2-phenylindole) and analyzed by epifluorescence microscopy (Axiophot II Imaging, Zeiss, Germany). Selected filters were analyzed by Confocal Laser Scanning Microscopy (LSM 780, Zeiss, Germany) including the Airyscan technology.

### Synthesis of authentic standards for metabolites

To produce alkyl-CoM standards, 1 g of coenzyme M was dissolved in 40 ml 30% (v/v) ammonium hydroxide solution and to this solution 1.8 to 2 g of bromoethane, bromoproane or bromobutane was added. The mixture was incubated for 5 h at room temperature under vigorous shaking and then acidified to pH 1 with HCl. The produced standard had a concentration of approx. 25 mg ml^−1^ which for mass spectrometry measurements was diluted to 10 μg ml^−1^.

### Extraction of metabolites from the Ethane50 culture

In the anoxic chamber 20 ml of Ethane50 culture was harvested into 50 ml centrifuge tubes. Tubes were centrifuged at 3000 rcf for 10 min and the supernatant was removed. The pellet was resuspended in 1 ml acetonitrile:methanol:water (4:4:2; v/v/v) mixture in lysing matrix tubes (MP Biomedicals, Eschwege, Germany) with glass beads. Afterwards the tubes were removed from the anoxic chamber and the samples were mechanical lysed in a FastPrep homogenizer (MP Bio) with 5 cycles with 6 M/s for 50 sec, and cooling on ice for 5 min in between the homogenization steps. Finally, the samples were centrifuged for 5 min at 13.000 × g and the supernatant transferred to a new tube and stored at −20°C.

### Solvents for LC-MS/MS

All organic solvents were LC-MS grade, using acetonitrile (ACN; BioSolve, Valkenswaard, The Netherlands), isopropanol (IPA; BioSolve, Valkenswaard, The Netherlands), and formic acid (FA; BioSolve, Valkenswaard, The Netherlands). Water was deionized by using the Astacus MembraPure system (MembraPure GmbH, Henningsdorf, Berlin, Germany).

### High resolution LC-MS/MS

The analysis was performed using a QExactive Plus Orbitrap (Thermo Fisher Scientific) equipped with an HESI probe and a Vanquish Horizon UHPLC System (Thermo Fisher Scientific). The metabolites from cell extracts were separated on an Accucore C30 column (150 × 2.1 mm, 2.6 μm, Thermo Fisher Scientific), at 40 °C, using a solvent gradient created from the mixture of the buffer A (5% Acetonitrile in water, 0.1% formic acid) and buffer B (90/10 IPA/ACN, 0.1% formic acid). The solvent gradient was the following: Fraction B (%) of 0, 0, 16, 45, 52, 58, 66, 70, 75, 97, 97, 15, 0, at −2 min. (pre run equilibration), 0, 2, 5.5, 9, 12, 14, 16, 18, 22, 25, 32.5, 33, 34.4 and 36 min. of each run, and a constant flow rate of 350 μl min^−1^. The samples injection volume was 10 μl. The MS measurements were acquired in negative mode for a mass detection range of 70–1000 Da. In alternation, a full MS and MS/MS scans of the eight most abundant precursor ions were acquired in negative mode. Dynamic exclusion was enabled for 30 seconds. The settings for full range MS1 were: mass resolution of 70,000 at 200 *m/z*, AGC target 5×10^5^, injection time 65 ms. Each MS1 was followed by MS2 scans with the settings: mass resolution 35,000 at 200 *m/z*, AGC target 1×10^6^, injection time 75 ms, loop count 8, isolation window 1 Da, collision energy was set to 30 eV.

### Determination of carbon back flux into the ethane pool

Aliquots of active AOM culture (50 ml) were transferred into 70 ml serum bottles with N_2_:CO_2_ headspace. In the SIP experiment addition of 99% ^13^C-labeled inorganic carbon (1 ml, 350 mM) led to δ ^13^C-DIC values of +25,000 ‰ as measured by cavity ringdown spectrometry. Ethane (2 atm = 1.8 mM) was added to both experiments and cultures were stored at 50°C. To determine the overall ethane oxidation activity sulfide concentrations were measured every few days as described above and converted to ethane oxidation rates using ratios of eq. 1. To measure the development of ethane δ^13^C values 1 ml of the gas phase was samples every few days, as stored it 10 ml Exetainer vials with 2 ml NaOH and measured ethane isotopic composition using gas chromatography coupled via a combustion oven to isotope ratio mass spectrometry (He as carrier gas, flow rate, column, temperature program).

### Net ethane production test

To test for net ethane production, in 156 ml serum flasks replicate incubations with about 0.5 g active Ethane50 culture (wet weight) in 100 sulfate-free medium were prepared. Four different conditions were tested in three biological replicates with the addition of (1) 1.5 atm H_2_; (2) replicate condition to (1) but only 0.05 g biomass (3) 1.5 atm H_2_ plus 28 mM sulfate (4) an activity control with addition of sulfate and 1.5 atm ethane. Cultures were incubated over 27 days at 50°C and sulfate and ethane concentrations were monitored as described above.

### Data availability

All sequence data are archived in the ENA database under the INSDC accession numbers PRJEB36446 and PRJEB36096. Sequence data from Loki’s Castle is archived under NCBI BioSample number SAMN13220465. The 16S rRNA gene amplicon reads have been submitted to the NCBI Sequence Read Archive (SRA) database under the accession number SRR8089822. All sequence information has been submitted using the data brokerage service of the German Federation for Biological Data (GFBio) [63], in compliance with the Minimal Information about any (X) Sequence (MIxS) standard [64], but some data is still under ENA embargo. For reviewers, sequence information is stored under https://owncloud.mpi-bremen.de/index.php/s/QSMycWOBB38AunL.

## Acknowledgements

We thank Susanne Menger for her contribution in culturing the target organisms, Janine Beckmann for metabolite analysis, and Dr. Heidi Taubner and Xavier Prieto Mollar (Hinrichs Lab, MARUM, University Bremen) for performing isotope analyses, We are indebted to Andreas Ellrott for assisting in confocal microscopy, and Gabriele Klockgether for her kind help with gas chromatography. We thank Matthew Schechter for analyzing the community compositions in a lab rotation. We also thank Tristan Wagner for vivid discussions on the metabolism of Ethanoperedens. We thank Bigelow SCGC for their work in sequencing single cell genomes used in this study. We are enormously grateful to I. Kostadinov and the GFBIO for the support and help during data submission. We are indebted to the crew and science party of R/V *Atlantis* and HOV *Alvin* Expedition AT37-06 (NSF Grant 1357238 to A. Teske). This study was funded by the Max Planck Society and the DFG Clusters of Excellence ‘The Ocean in the Earth System’ and ‘The Ocean Floor – Earth’s Uncharted Interface’ at MARUM, University Bremen. Additional funds came from the ERC ABYSS (Grant Agreement No. 294757) to A.B..

## Author contributions

C.H., K.K. and G.W. designed the research. A.T. and G.W. retrieved the original Guaymas Basin sediment sample. F.V., K.-M., V., R. S., I.H.S. and A.B. retrieved additional samples. C.H. and G.W. performed the cultivation, physiology and isotope experiments. C.H., F.V. K.-M. V. and K.K. performed fluorescence microscopy. C.H., M.L. and G.W. performed metabolite analysis. C.H., R. L.-P. M., R.A. K.K. and G.W. performed metagenomic and phylogenetic analyses and developed the metabolic model, C.H., and G.W. wrote the manuscript with contributions from all co-authors.

---

**Table S1** PCR primers used the amplification of archaeal and bacteria 16S rRNA genes and oligonucleotide probes used for CARD-FISH.

**Table S2** Summary of single cell and metagenome assembled genomes presented in this study and average nucleotide and amino acid identities. ANI and AAI values were calculated with publically available genomes and genomes presented in this study. Enveomics tools were used for the calculation [50].

**Table S3** Genomes and gene expression data of the Ethane50 culture and overview of genes potentially involved in the ethane metabolism and electron cycling in the Ethane50 culture. Expression values shown in triplicates for *Ca*. E. thermophilum and *Ca*. D. auxilii.

**Table S4** Overview of environmental sampling sites used for this study.

**Table S5** Marker genes used for calculation of genome tree based on archaeal marker genes presented in Rinke, Schwientek [58].

**Table S6** Summary dataset of the development of substrates and products in the Ethane50 enrichment culture. Development of ethane, sulfide and sulfate concentrations in E50 culture in triplicates. Development of sulfide concentration in E50 culture in 10 replicates. Development of ethane and sulfide concentrations in triplicates of the Ethane50 culture with hydrogen gas (1.5 atm) with and without sulfate. The positive control contained 1.5 bar ethane and sulfate.

**Fig. S1.**
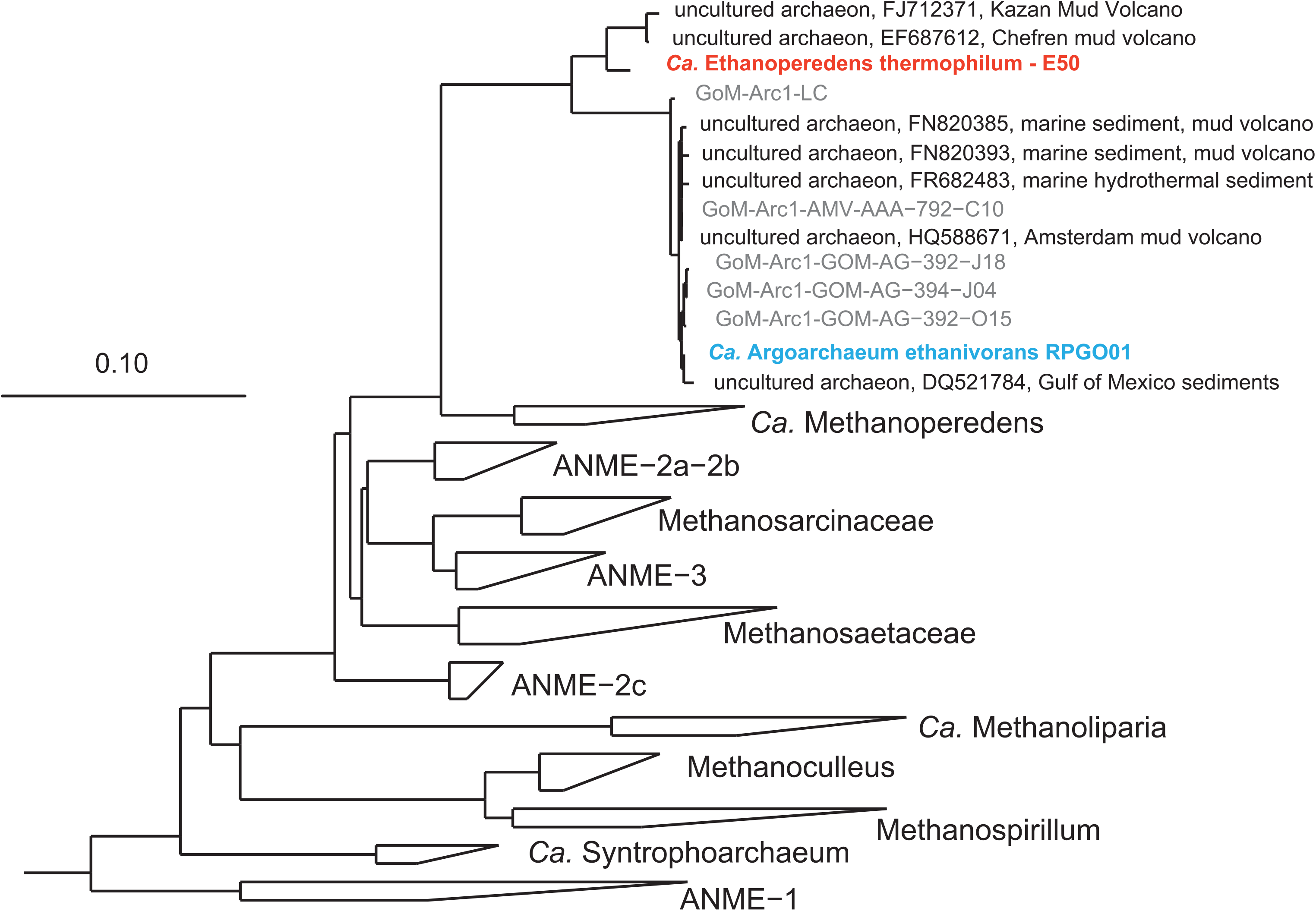
Phylogenetic affiliation of the GoM-Arc1 clade archaea with other archaea based on 16S rRNA gene comparison. The tree was constructed using ARB [56] and the FastTree 2 package [57] using a 50% similarity filter. 410 sequences with a length of at least 1100 bp, excluding partial sequences retrieved from single cells, were used. Bar shows 10% sequence divergence.

**Fig. S2.**
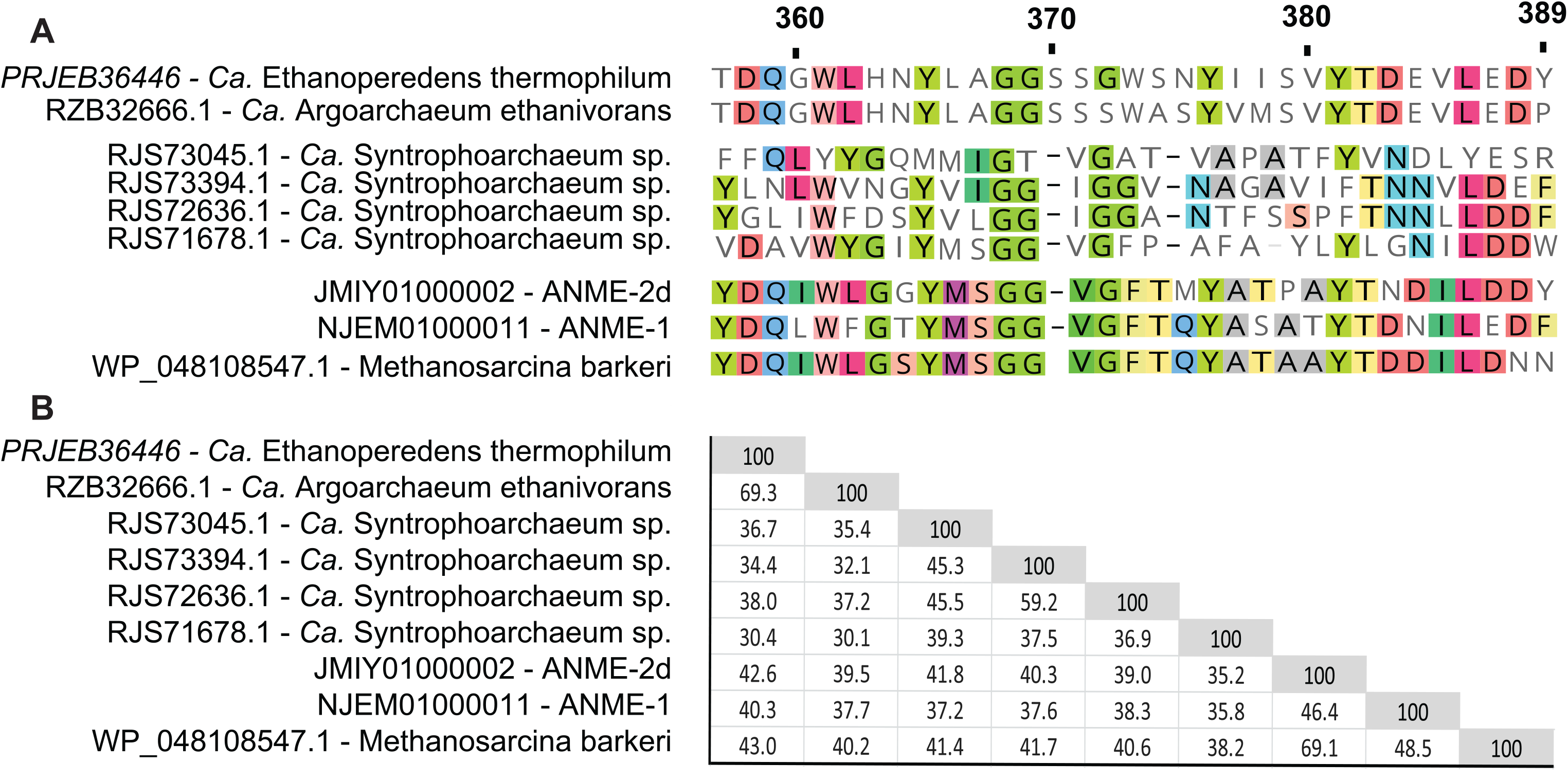
Comparison of *mcr*A sequences from the GoM-Arc1 clade to described canonical and non-canonical *mcr*A sequences. A, alignment of the active site of the *mcr*A from different representative genomes. The four different *Ca*. Syntrophoarchaeum sequences belong to the same genome bin. Amino acid positions refer to *Ca*. E. thermophilum E50 mcrA sequence. B, identity matrix of *mcr*A sequences based on NCBI blastp alignment.

**Fig. S3.**
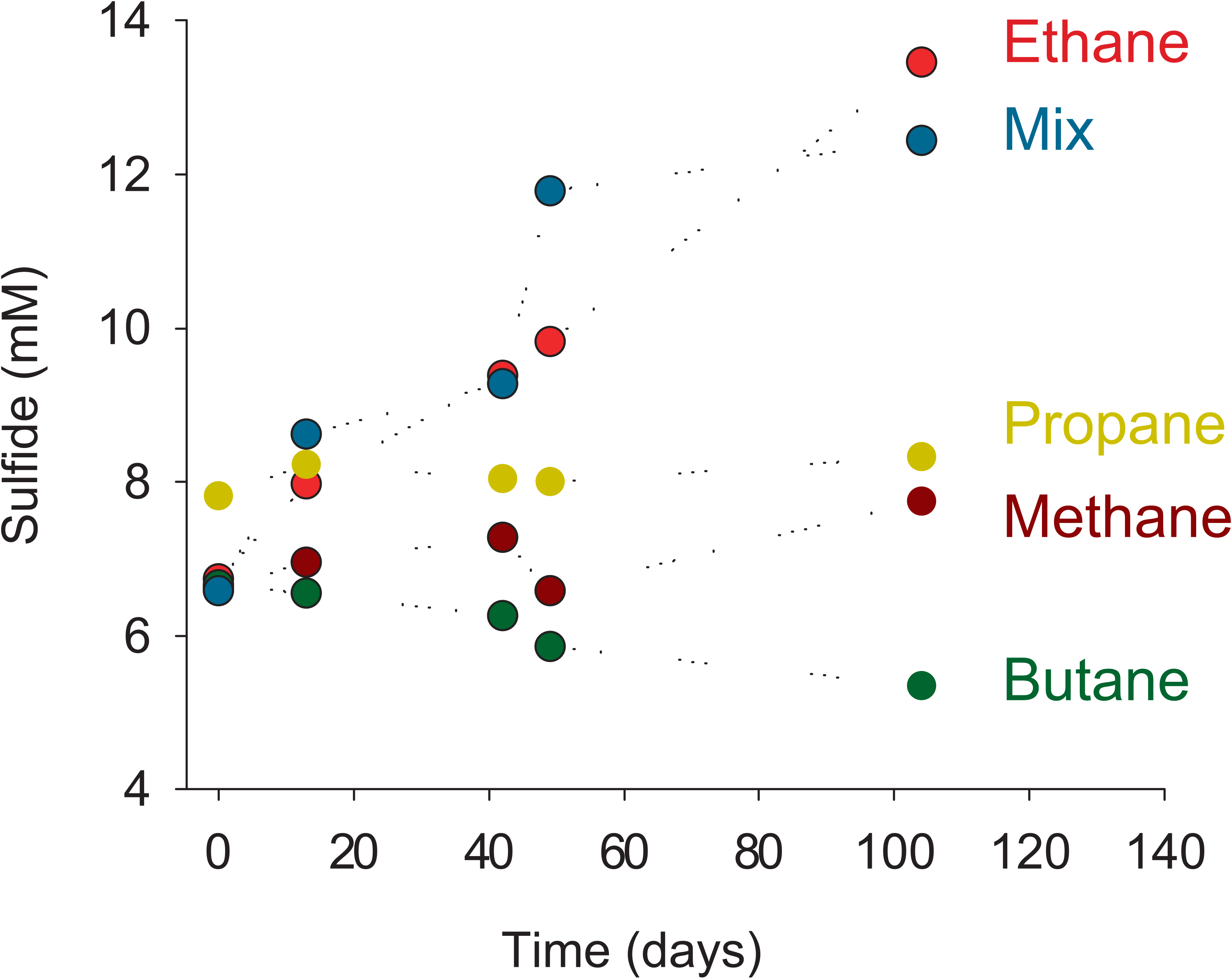
Development of sulfide concentrations in substrate experiments in replicate incubations of the Ethane50 culture supplied with (**A**) ethane, alternative alkanes or a mix of these substrates; and (**B**) ethane and sulfate compared to ethane with elemental sulfur or only elemental sulfur. Results show that that ethane is the only alkane used as electron donor in Ca. E. thermophilum, and sulfate is the only used electron acceptor. Further, elemental sulfur is not disproportionated.

**Fig. S4.**
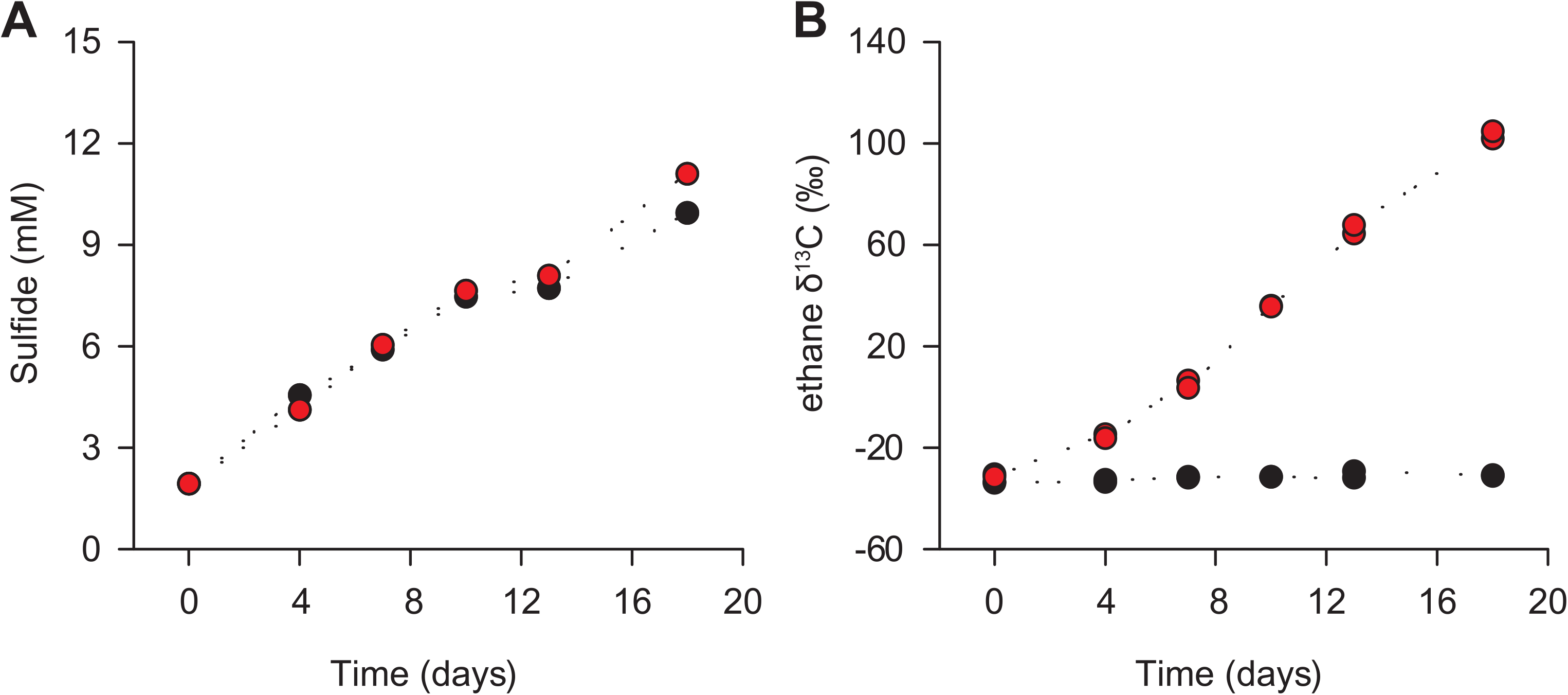
Test for the transfer of inorganic carbon into ethane in the ethane 50 culture. **A,** development of sulfide concentrations in the culture with ethane as energy source and sulfate as electron acceptor **B,** development of δ^13^C values in ethane in the two cultures (controls δ13C_DIC_ −35‰) ^13^C-DIC amended culture with δ^13^C_DIC_ =+25994 ‰. Based on simple mass balance calculations on the development of fractions we infer that sulfate-dependent anaerobic AOM in these enrichments is accompanied by a back flow of inorganic carbon amounting to 1-3% of the forward rate. This back reaction indicates a general reversibility of ethane oxidation.

